# Hippocampal reconfiguration of events in mnemonic networks

**DOI:** 10.1101/2021.05.25.445607

**Authors:** Silvy H.P. Collin, Branka Milivojevic, Christian F. Doeller

## Abstract

It is widely assumed that episodic memories are not stored in isolation but rather in dynamic event networks. However, the mechanisms of the underlying dynamic of these representations, in particular how such networks are updated, remain elusive. In this study, we investigated the reconfiguration of events into event networks in the hippocampus by presenting new events that could update either one of two competing narratives. During the first session, participants viewed four animated movies, each representing a distinct narrative; two distinct narratives from the Jones family and two distinct narratives from the Smith family. During the second session, we re-exposed participants to snapshots of these narratives along with snapshots of new events from one of the two families, allowing updating of the acquired event networks of that family. Our findings show that the hippocampus integrated new events that relate to the old family, and then integrated these new events with the corresponding old events. Furthermore, hippocampal representations of the events within a narrative became better integrated after updating. Our results shed new light on the neural mechanisms underlying flexible mnemonic updating with realistic events and further advance our understanding of the structured reconfiguration of event networks in memory.

## INTRODUCTION

Episodic memories, which are stored as networks of interrelated events (Eichenbaum et al., 1999; Shohamy and Wagner, 2008; Zeithamova et al., 2012a; Schapiro et al., 2013; Ezzyat and Davachi, 2014; Collin et al., 2015; Milivojevic et al., 2015; Backus et al., 2016; Deuker et al., 2016; Schlichting and Frankland, 2017), are remarkably malleable. We can integrate new experiences with pre-existing memories, even if it requires reinterpretation of prior knowledge, a function which involves hippocampus, medial prefrontal and orbitofrontal cortices (Shohamy and Wagner, 2008; Staresina and Davachi, 2009; Zeithamova et al., 2012b; Shohamy and Turk-Browne, 2013; Schlichting et al., 2015; Kitamura et al., 2017; Zhou et al., 2021). Updating of memories (Forcato et al., 2007; Hupbach et al., 2007, 2008; Schiller et al., 2010; Newman and Norman, 2010; Schiller and Phelps, 2011; Kuhl et al., 2012; Jacques and Schacter, 2013; Kroes et al., 2014; Poppenk and Norman, 2014; Schlichting and Preston, 2016; Richter et al., 2019) can trigger reconsolidation (Misanin et al., 1968; Nadel and Land, 2000; Nader et al., 2000; Wiltgen et al., 2004; Lee et al., 2017) whereby consolidated memories can become susceptible to change after reactivation. However, the neural mechanisms underlying episodic memory updating remain elusive.

Piaget (Piaget, 1929) suggested that we remember by using mental structures of related information or schemas (Bartlett, 1932). He further proposed that updating of schemas would be possible either by modifying new information to fit the schema, or restructuring the schema to accommodate new information (Piaget, 1929; Preston and Eichenbaum, 2013). Several studies have investigated neural mechanisms of schemas and linked them to hippocampus and frontal cortex involvement (Bartlett, 1932; Tse et al., 2007, 2011; Van Kesteren et al., 2012, 2010; Zeithamova et al., 2012a; Kumaran, 2013; McClelland, 2013; Preston and Eichenbaum, 2013; van Buuren et al., 2014; Baldassano et al., 2018; Raykov et al., 2020; Zhou et al., 2021). However, most studies investigated schemas by using relatively simple forms of item-item associations or by investigating schema congruency based on prior semantic knowledge. (Van Kesteren et al., 2010) and (Baldassano et al., 2018) have investigated neural mechanisms of schema formation in a more realistic setting, focusing on hippocampal-mPFC connectivity and posterior medial and frontal cortices respectively. Nevertheless, it remains unknown how the hippocampus is involved.

Research suggests that the hippocampus is involved in representing event networks (McKenzie et al., 2014; Collin et al., 2015, 2017; Milivojevic et al., 2015, 2016; Bonasia et al., 2018; Brunec et al., 2018; Guo and Yang, 2020; Chang et al., 2021). It remains unclear what the hippocampal mechanisms are for updating such networks. We used pattern-similarity analysis of fMRI-data to examine the representational geometry of events in the hippocampus during and after updating. We hypothesized that integration of new events into pre-existing event networks would be similar to restructuring hippocampal schemas reported in rodents (McKenzie et al., 2013) and should stimulate updating based on modification of pre-existing hippocampal representations of event networks (Collin et al., 2015; Milivojevic et al., 2015, 2016; Brunec et al., 2018).

Participants were exposed to four narratives (each consisting of multiple events) on Day-1 of which the two narratives featuring the same family were updated with new events on Day-2. Thus, the stimuli had a hierarchical structure (“Family – Day – Event”) which allowed us to investigate episodic memory updating at various levels. Participants were asked to actively place each new event at a specific timepoint into one narratives which forced them to re-organize that narrative. Before and after being presented with new events, participants were presented with old events, which allowed us to measure the degree of reorganization of the networks as a consequence of updating. Firstly, we predicted that the different events within a narrative would become better integrated due to updating compared to those narratives where no updating occurred. Secondly, we predicted that updating would lead to better integration of all events from the same family, also across the different narratives from the same family. Thirdly, we predicted more stable event-specific representations due to updating.

## MATERIALS & METHODS

### Participants

Thirty-four students participated in this experiment. All participants were right-handed. They were recruited via an online participant recruitment system from Radboud University. Four participants were excluded from further analyses due to excessive head motion (2), or due to incomplete data sets (2). The final group consisted of 30 participants (11 males, aged 18–44 years, and mean age 24.3) who all had normal or corrected-to-normal vision. The experiment was approved by the local ethical review committee (CMO region Arnhem-Nijmegen, NL) and participants gave written informed consent to participate.

### Experimental design

#### Stimulus material

Stimuli consisted of animated cartoon narratives generated using The Sims 3 (www.thesims3.com) life-simulation game. With this game, we created several characters who were grouped into two virtual families (Smith Family and Jones Family) with three family members each. In this experiment, participants were presented with four narrative-videos (i.e. animated cartoon narratives) in total (two per family). Each narrative-video had a duration of four minutes. Each of these four-minute narratives comprised 10 events on average (See methods for more details) that together formed a typical day of that family. Full descriptions of the four narratives are available in the methods section. An additional behavioral experiment showed that the four narratives used were of similar complexity (see additional behavioral experiment 1 for more details). For the tasks in the MRI scanner (on Day 2 of the experiment), we used snapshots of the videos that were equalized for a number of image properties: color (images were gray-scale) and luminance, using the SHINE toolbox (Willenbockel et al., 2010) with the aim of minimising potential low level visual confounds in the analyses of the MRI data.

#### Day 1

On Day 1 of the experiment, participants underwent a ‘learning phase’ during which they watched the four narratives (three repetitions per narrative, the first narrative was only repeated after all narratives were seen once). Subsequently, they completed a free recall task during which they were asked to recall what happened in the four narratives.

#### Day 2

On Day 2 of the experiment, participants were presented with five task blocks (see Figure 1, note that task blocks [2], [3] and [4] were shown in the MRI scanner): [1] pre-scanning event-configuration memory test, [2] pre-block (old events were shown), [3] updating-block (new events were shown), [4] post-block (old events were shown again) and [5] post-scanning event-configuration memory test.

**Fig. 1.**
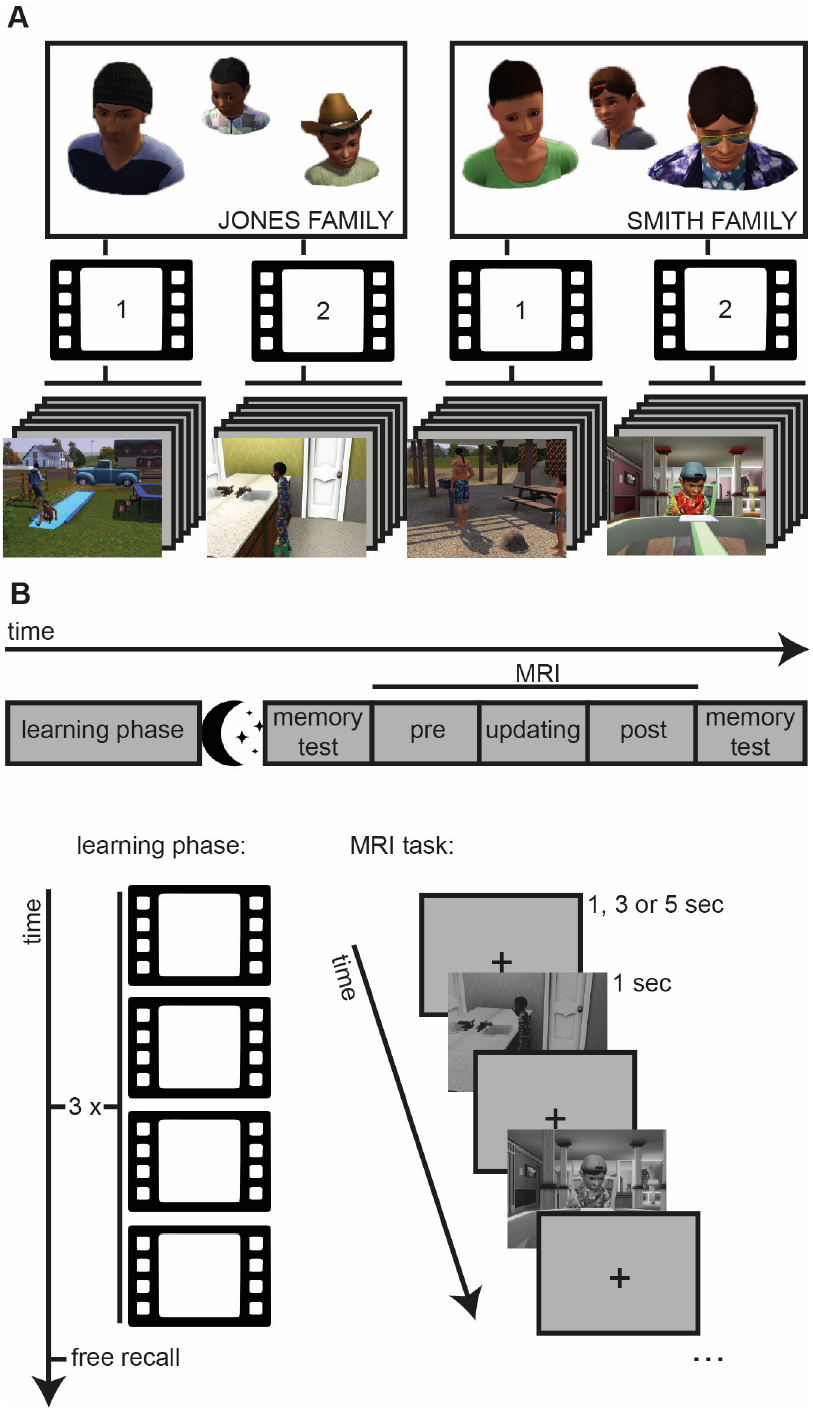
Experimental design. **(A) Hierarchical components of the narratives**. Illustration of two virtual families created using TheSims3 life-simulation game. Participants (N=30) were presented with four narratives in total (two per family). Each of these four, four-minute narratives comprised a number of events that together formed a typical day of one of the families. Full descriptions of the four narratives are available in the methods. A behavioral experiment in which participants had to segment the narratives into different events showed that a similar number of events could be recognized for the four narratives (see methods for details). **(B) Overview of the experimental sessions**. The experiment started with a ‘learning phase’ during which participants watched the four narratives (three times in pseudorandom order) and completed a free recall task. The next day included five subsequent task blocks in the following order: memory test, pre block with re-exposure to events from the original narratives, updating block where new events were introduced for one of the families (importantly, only the narratives of one family were updated with new events) along with Control Events, post block with re-exposure to events from the original narratives, and a second memory test. These memory tests included event-by-event memory judgments of how the events are related. The new events were deliberately designed in such a way to be neutrally related to both narratives per family, as to allow for subject-specific flexible recombination with either narrative of the corresponding family (methods for more details).

#### Pre-scanning event-configuration memory test

During the first task block ([1] event-configuration memory test), participants were presented with 24 snapshots of the narratives (six per narrative) and were told to arrange these snapshots in a circular arena based on how these snapshots belong together (see (Charest et al., 2014)). They performed this task for ten minutes.

#### Pre block

During the second task block ([2] Pre-block) participants were presented with these 24 snapshots of narrative-events in the MRI scanner, one at a time, in a pseudorandom order (twelve repetitions per snapshot; 1 sec stimulus duration; 1, 3 or 5 sec intertrial interval). The trial order was pseudorandom to guarantee a roughly equal distribution of the 24 stimuli over time. This was done by dividing the entire task in twelve subsequently presented ‘sub-blocks’. In each of these ‘sub-blocks’, all 24 snapshots were presented once in a random order. Each of the ‘sub-blocks’ contained an equal number of 1, 3 and 5 sec intertrial intervals.

#### Updating block

During the third task block ([3] Updating-block) participants were presented with twelve new events in the MRI scanner, one at a time, in a pseudorandom order (twelve repetitions per snapshot; 1 sec stimulus duration; 1, 3 or 5 sec intertrial interval). In this task block the trial order was pseudorandom and via the same logic as in the second task block (see above). Although all of these twelve new events showed a completely new event, six of them featured one of the original virtual families (New Narrative Events) while the other six new events featured a new set of people (New Control Events). Thus, only the narratives of one virtual family was updated with new events (which narratives were updated was counterbalanced across participants). Participants received the following instruction for this task block: “You will now see a number of new events. It is your task to figure out whether a new event contains people you have already seen, or new people. If the event contains new people, it is a completely new event. However, if the event contains people you have already seen, then this event belongs to one of the narratives you have already seen earlier. In this case, it is your task to figure out to which narrative exactly this event belongs.”

#### Post block

During the fourth task block ([4] Post-block) participants were again presented with the 24 snapshots of the original narrative-events in the MRI scanner, one at a time, in a pseudorandom order (twelve repetitions per snapshot; 1 sec stimulus duration; 1, 3 or 5 sec intertrial interval). The trial order and intertrial intervals in this task block were exactly the same as in the pre-block (i.e. task block 2) to ensure the same timing between all stimulus presentations in these two task blocks.

#### Target-detection task

Additionally, during above described task blocks two, three and four, participants performed a target detection task to ensure continuous attention of participants throughout the task. Participants were told to press button ‘1’ when they saw a snapshot of a girl on a scooter (10 % of the total number of stimuli randomly distributed throughout the entire task block, this target stimulus was not related to any other stimulus), and button ‘2’ when they saw any other stimulus.

#### Post-scanning event-configuration memory test

During the fifth task block ([5] event-configuration Memory test) participants completed a memory test which was similar to the pre-scanning memory test, but now they were presented with all 36 snapshots (24 original events and 12 new events). They again had to arrange these snapshots in a circular arena based on how these snapshots belong together. They performed this task for fifteen minutes (total duration longer to keep the average time per snapshot the same for both memory tests).

##### Full descriptions of the four narratives and of the new events introduced on Day 2

###### Jones Family – Weekend narrative

This narrative describes a weekend day of dad, son, and daughter Jones. It starts with the kids playing on a seesaw in the garden of their home. <u>Then they go to their dad and talk to him</u> for a while. They walk away and get ice-cream from an ice-cream truck in front of their home. <u>They eat the ice-cream</u>, and after that play again in their garden, <u>on the seesaw</u> and in the sandpit. Dad joins the kids after a while. Then the camera switches to the other side of the garden. First the daughter and later also dad and son Jones enter in their swimming clothes, and <u>play with water</u> for a while. When it starts to get darker, the kids, still in swimming clothes, play for a little while in the sandpit again, but then Dad comes to tell them to go inside. <u>Dad goes into the bathroom in his swimming clothes</u>, and leaves again, dressed. The daughter goes into the bathroom wearing her swimming clothes, and you see her fill the bath. In the next scene, dad and son are in the kitchen, and soon the daughter also joins. <u>They eat pizza</u> together at the kitchen table. Then, dad washes the dishes.

###### Jones Family – Weekday narrative

This narrative describes a weekday of dad, son, and daughter Jones. The kids get out of bed. Meanwhile, <u>dad is preparing breakfast</u>. <u>The kids have a pillow fight</u>. After a while, <u>the kids enter the kitchen where dad is still preparing breakfast</u>. They have breakfast together at the kitchen table, and talk a bit with each other while having breakfast. Then, <u>the daughter enters the bathroom</u> and brushes her teeth, and subsequently the son enters the bathroom and <u>he brushes his teeth</u> too. They walk out of the front door, and <u>get in the school bus</u> which is parked in front of their home. The bus drives them to school, meanwhile passing another car, a man who is jogging, a man with a horse, and a couple of buildings. The last scene shows that the kids get out of the school bus and walk into the front door of the school.

##### Smith family – Weekend narrative

This narrative describes a weekend day of mum, dad, and son Smith. In the first scene, <u>they are all in the kitchen preparing food</u>. Later, you see mum also eating some food. Dad and son then leave the kitchen. The <u>son goes into his bedroom and cleans up a bit</u>. The father gets in his car and drives to the beach. At the beach, he goes into the water to swim. In the next scene, <u>the son asks mum something</u>. She agrees to what he is asking, which makes the son cheer. Mum prepares some sausages in the kitchen. Then she goes to the son and says goodbye to him. The son gets on his bike and <u>bikes to the beach</u> as well. He passes a man on a horse. Once he is at the beach, <u>he joins his dad with swimming</u>. After they have been swimming for a while, they go onto the beach, and <u>dad prepares some sausages for him and his son on a barbecue</u>. His phone rings and he talks in his phone for a while.

###### Smith family – Weekday narrative

This narrative describes a weekday of mum, dad, and son Smith. Mum and dad are jogging on the sidewalk in front of their home. <u>Once they reach their home, they enter via the front door</u>. In the next scene, the son leaves school and gets into the school bus. The school bus drives through the town for a while. Then, the <u>son arrives at home</u> and enters the house via the back door. <u>Mum greets him</u>, and then tells him something. The son walks to the kitchen table to <u>do some homework</u>. Meanwhile, mum is preparing food in the kitchen. She uses a blender and the oven. Once the food is ready, mum, dad, and son eat together at the kitchen table. After dinner, the son watches TV. After a while, dad comes to talk to him. Subsequently, the son leaves to the bathroom and <u>brushes his teeth</u>. Then he goes into his bedroom, and goes to bed. Mum and dad are on the couch together, talk for a while and <u>watch TV together</u>.

*(Please note that the underlined text parts in these descriptions are the scenes that were shown in the MRI scanner during the pre and post blocks)*

###### Jones Family – New events

The six new events of the Jones family include:

1. Daughter reading a book in the living room
2. Dad reading the newspaper
3. Dad drinking a soda in the kitchen
4. The kids watching TV
5. Son throwing garbage into the bin
6. Dad in front of the house with a dog

###### Smith Family – New events

The six new events of the Smith family include:

1. Dad making a cup of coffee
2. Son doing some exercises in front of the TV
3. Son playing with a parrot
4. Mum and dad in a home office with dad in front of a computer
5. Son painting on a canvas
6. Mum, dad and son sitting on the porch outside their home

###### Control events

The six control events each feature a different character doing a different activity:

1. A woman at a football table
2. A man in a kitchen on a laptop
3. A man and woman in the gym
4. A woman sitting outside on a bench
5. A man and woman in a living room
6. A woman sunbathing in her garden

### Image acquisition

Imaging data were acquired on a 3T Siemens Prisma scanner using a 32-channel head coil. We used a 2D echo-planar imaging (EPI) sequence with the following parameters: volume TR = 2000 ms; time echo (TE) = 24 ms; flip angle = 85°; volume resolution = 2 mm^3^; field of view (FOV) = 210×210×74 mm; acceleration factor = 3. Since we used a reduced FOV (i.e., 37 slices), we acquired an AutoAlign Head LS scan for automated positioning and alignment using anatomical landmarks (to ensure that the same anatomical part of the brain was scanned in all subjects). The structural T1-weighted image was acquired using an MPRAGE-grappa sequence with the following parameters: TR = 2300 ms; TE = 3.03 ms; flip angle = 8°; in-plane resolution = 256×256 mm; number of slices = 192; acceleration factor PE = 2; voxel resolution = 1 mm^3^, duration = 321 s.

### Image preprocessing

Image preprocessing was performed using the Automatic Analysis Toolbox (Cusack et al., 2015), which uses custom scripts combined with core functions from SPM8 (www.fil.ion.ucl.ac.uk/spm), FreeSurfer (http://surfer.nmr.mgh.harvard.edu/) and FSL (http://fsl.fmrib.ox.ac.uk/fsl/fslwiki/). We used the SPM8 functional image realignment procedure to estimate movement parameters (three for rotation and three for translation). To improve the structural image quality, we bias-corrected the structural image (Ashburner and Friston, 2005) and de-noised it using an optimised non-local means filter (Manjón et al., 2010). Subsequently, the functional images were co-registered to the structural image using a four-step procedure: 1) the structural image was co-registered to the T1 template; 2) the mean EPI was co-registered to the EPI template; 3) the co-registered mean EPI was co-registered to the structural image; 4) the orientation parameters of the mean EPI were applied to the individual EPIs. For each structural and mean EPI, we used the FSL brain extraction toolbox (Smith, 2002) to create a structural and functional brain-only mask. The resulting skull-stripped structural image was segmented into grey matter (GM), white matter (WM) and cerebro-spinal fluid (CSF) (Ashburner and Friston, 2005). During first-level GLM modelling, the six movement parameters were used as nuisance regressors. For the multivariate analyses (see below), the images were not pre-processed further. For the univariate (control) analysis, the functional images were normalized to the MNI template using normalization parameters estimated through the unified segmentation procedures, as implemented in SPM8 (Ashburner and Friston, 2005), and smoothed using an eight mm full-width at half maximum (FWHM) 3D Gaussian kernel.

### Representational similarity analysis

We used representational similarity analysis (RSA) to analyse the multivoxel pattern of neural activity and applied a roving searchlight approach on our whole-brain data using version 3 of the RSA toolbox developed by Kriegeskorte et al (Cognition and Brain Science Unit, Cambridge). To this end, we examined the Spearman correlation coefficients between patterns of activity within spherical regions of interest (ROIs), or search spheres, throughout the whole brain volume.

First-level modelling was performed using a modified version of Mumford, Turner, Ashby and Poldrack (Mumford et al., 2012) approach whereby estimate of each regressor of interest was estimated by running a separate GLM with two regressors, one being the regressor interest, and the other modelling all other trials. We performed 60 separate GLMs corresponding to the 60 regressors of interest. This included one regressor for each of the 24 events in the pre-block, one regressor for each of the twelve events in the updating-block, and one regressor for each of the 24 events in the post-block. Each of these regressors of interest modelled twelve trials. To investigate the lowest level of the hierarchy (i.e. event representation), we split each regressor into two regressors, one modeling the odd events, and one modeling the even events. Each model also included the following as part of a nuisance regressor: targets and button responses (from the target-detection task). All regressors of no interest were convolved with the canonical hemodynamic response function, producing a modelled time-course of neural activity. Additionally, six nuisance regressors per imaging run were included to control for head movement. Voxel-wise beta estimates resulting from the 60 regressors of interest were used for the subsequent searchlight RSA.

In the second analysis step, we investigated the degree of correlation between patterns of activity within search spheres measuring 12 mm in diameter, moving this search sphere across the entire field of view. Each hypothesis regarding the change in neural similarity was then evaluated in each participant using a model representational dissimilarity matrix (RDM), and comparing this model RDM with an RDM of each search sphere. The searchlight analysis was performed on the native space images of each participant by moving the centre of the search sphere through the grey-matter masked volume one voxel at a time. Resulting single-subject statistics were mapped back to the centre voxel of each spherical ROI, thus yielding a single-subject neural-similarity map that was entered into a group analysis. The first-level results were then normalized to the MNI template using normalization parameters estimated through the unified segmentation procedure of SPM8, and a smoothing Gaussian kernel of 8 mm^3^ FWHM was applied to these data. A second-level model was carried out to examine changes in neural similarity at the group level. Note that the peak voxels represent neural similarity values across the grey-matter voxels within the spherical searchlights centred on the peak voxels.

### Statistical analyses (representational similarity analysis)

At the second-level, a one-sample T test (with 5000 permutations) using the Statistical NonParametric Mapping toolbox was used (SnPM13, http://warwick.ac.uk/snpm). A small-volume correction was performed to test for statistical significance of the RSA effects in the hippocampus (i.e. small-volume correction with an ROI including the left and right hippocampus based on the AAL template was used). Brain maps included in figures are presented in neurological convention.

RSA prediction matrices of analysis 2: To test for representation of the new events in the brain, we used a prediction matrix that predicted high similarity between the six new events, and low similarity between the six control events. Additional control analysis: To investigate whether this effect could be driven by higher overlap between events (i.e., the same characters appearing in multiple new events, but in contrast to that, a different character appearing in each of the control events), we used prediction matrices for new events according to common people within the events (i.e. this control analysis would test whether increased similarity between new events was partly driven by common characters appearing in those new events). Thus, this analysis only included comparisons between new events (excluding the six control events), and the prediction matrices reflected high similarity between new events in which the same character was present and low similarity between new events in which different characters were present).

RSA Prediction matrices of analysis 3: To test for representations of the three hierarchical levels, we tested 3 prediction matrices for which we used: [1] a contrast that predicted higher neural similarity within compared to across events ignoring all comparisons across narratives/families (event representation), [2] a contrast that predicted higher neural similarity for within-narrative event pairs compared to across-narrative event pairs ignoring all comparisons across family (narrative representation), and [3] a contrast that predicted higher neural similarity for within-family event pairs compared to across-family event pairs ignoring all comparisons within-narrative (family representation). We then tested with a repeated-measures ANOVA whether these hierarchical levels were present in the brain.

RSA Prediction matrices of Analysis 4: To test for integration of New Narrative Events with Old Narrative Events, we used a prediction matrix that predicted high similarity between New Narrative Events with Old Narrative Events from the updated family, and low similarity between New Narrative Events with Old Narrative Events from the non-updated family as well as low similarity between Control Events and Narrative Events from the updated and non-updated family. This included presentations of Narrative Events pre and post updating. We repeated this analysis for Narrative Event pre updating and Narrative Events post updating separately.

### Univariate control analysis

Normalised and smooth data were used for the univariate control analysis implemented in SPM8. The purpose of this analysis was to examine whether any differences were evident in univariate signal amplitude, which may confound RSA results. Therefore, we used the same first-level model for both univariate and RSA analyses. This included one regressor for each of the 24 events in the pre-block, one regressor for each of the twelve events in the updating-block, and one regressor for each of the 24 events in the post-block, and one regressor for targets and one for button responses. Additionally, we included six nuisance regressors per imaging run to control for head movement. First-level contrasts were generated to test for potential amplitude differences between the events. The following T-contrasts were performed: higher activity during Narrative Events of the updated family compared to the non-updated family, and higher activity during New Events relative to Control Events. These T-contrasts were tested at a group level using second-level modelling in SPM8. At a liberal threshold of P < .001 uncorrected, neither of these contrasts showed a significant effect in the hippocampus. Additionally, we extracted the mean amplitude for the exact location within the hippocampus where we found our main effects, which we tested with one-sample T-tests for statistical significance. There was no difference in (univariate) amplitude of the MR signal in the posterior hippocampus for updated vs non-updated family narrative events (two-tailed one-sample T-test, T(1,29) = 0.969, P = 0.34), neither for new vs control events (two-tailed one-sample T-test, T(1,29)= - 0.207, P = 0.837). There was no difference in (univariate) amplitude of the MR signal in the anterior hippocampus for updated vs non-updated family narrative events (two-tailed one-sample T-test, T(1,29) = - 1.348, P = 0.189), neither for new vs control events (two-tailed one-sample T-test, T(1,29) = 1.294, P = 0.206).

### Low level visual features control analysis

The snapshots of the videos used in the MRI blocks of the experiment were equalized on a number of low level visual features as described above in the ‘task design – stimulus material’ section of the methods. Nevertheless, to further rule out that any of the reported effects are driven by low level visual similarity, we computed visual similarity measures between all snapshots used in the experiment. For each snapshot, we computed a 2-dimensional discrete Fast Fourier Transform (FFT). From this we derived the imaginary and real parts of the FFT and hence calculated the magnitude and phase metrics. These metrics were then vectorised and correlated between all events. This lead to two matrices, one for FFT magnitude and one for FFT angle, which were then used as model RDMs in a new RSA analysis of the across-voxel patterns (i.e. an fMRI searchlight RSA). Again, we investigated these two low-level visual features dimensions at the exact location where we found our main effects, which we tested with repeated measures ANOVAs for statistical significance. There was no significant effect of low-level visual features in the posterior hippocampus (two-tailed paired-samples T-test: T(1,29) = 1.554, P = 0.131) or anterior hippocampus (two-tailed paired-samples T-test: T(1,29) = 0.447, P = 0.658).

### Statistical analyses (behavior)

To test for family and narrative hierarchical-level-based grouping effects in the behavioral results of the pre-scanning and post-scanning memory tests, we correlated both the family prediction matrix (i.e. high similarity within-family, low similarity across-family, excluding the within-narrative pairs) and narrative prediction matrix (i.e. high similarity within-narrative, low similarity across-narrative, excluding the diagonal) with the result matrix from the memory test for each participant separately (which included event-by-event distance measures). We then tested the mean correlation, separately for pre and post-scanning memory tests, and separately for family and narrative effect, in two-tailed one-sample T-tests, in which we used the threshold for statistical significance of P < 0.00625 (i.e. P < 0.05 corrected for multiple comparisons).

### Behavioral control experiments used during development of the task

#### Control experiment 1: Narrative-segmentation task

We aimed to determine if the complexity of the narratives (four minutes in duration each) used for the main experiment was indeed similar across the four narratives. We relied on the ability of observers to segment everyday activities into meaningful events (Zacks and Swallow, 2007).

Sixteen participants (five male, aged 18 to 29 years, and mean age 23.2) participated in this behavioral control experiment (referred to as ‘Narrative Segmentation Task’). They were recruited via an online participant recruitment system from the Radboud University. They all had normal or corrected-to-normal vision. The experiment was approved by the local ethical review committee (CMO region Arnhem-Nijmegen, NL), and participants gave written informed consent to participate. We used the same task procedure as described in prior studies investigating event segmentation and/or event boundaries (Zacks et al., 2001; Kurby and Zacks, 2008; Swallow et al., 2009) including three task blocks: [1] Passive viewing task, [2] Fine segmentation task, [3] Coarse segmentation task. In the passive viewing task, participants watched the four narratives (four minutes each) that are also used in the main experiment in a random order. The participants simply watched each video passively and were instructed to learn as much about the movie as possible. The task started with a practice video (1.5-minute duration). This task block was included to familiarize participants with the narratives. In the fine segmentation task, participants watched the same videos as those in the passive viewing task (again in a random order), but were asked to press a button at the points at which they believed one meaning and natural unit of activity ended and another began. This procedure has been shown to reliably measure the perceptual units of ongoing behavior. The participants were instructed to identify the smallest units of activity that seemed natural and meaningful. This task block also started with performing the same task on the 1.5-minute duration practice video. After the practice video, participants were asked if the task was clear, after which they proceeded with the actual videos. The coarse segmentation task was identical to the fine segmentation task, except that the participants were asked to identify the *largest* units that were natural and meaningful to them. This task block also started with performing the same task on the 1.5-minute duration practice video. After the practice video, participants were asked if the task was clear, after which they proceeded with the actual videos. The order of the fine and coarse segmentation task (i.e. task block two and three) was counterbalanced across participants. Separately for the coarse and the fine segmentation of the four videos, we calculated the number of button presses during the presentation of the video (separately for each of the four videos). The button responses correspond to a perceived event boundary. These values were added to a 2×2 repeated measures ANOVA with Segmentation (Coarse, Fine) and Narrative (1, 2, 3, and 4) as within-subject factors. This analysis showed that there are no differences in narrative complexity between the four narratives (F (3,45) = 1.157, P = 0.337).

#### Control experiment 2: Consistency task

To examine the consistency of the new events with the original narratives, we conducted a behavioral control experiment. In this control experiment (fifteen participants, one male, aged 19 to 28 years, mean age 22.1, recruited via online participant recruitment system of Radboud University, experiment approved by local ethical review committee, participants gave written informed consent) snapshots from the original narrative were presented on top of the computer screen and one of the new events on the bottom of the screen. All snapshots used in the main experiment were included in this control experiment in a random order.

Participants had to answer the question: ‘How consistent is the image below with the narrative above?’ They answered on a 6-point scale from not consistent (−3) to consistent (3). The task was self-paced. There were trials for each new event relative to each of the narratives. Average consistency of the new events to their original narratives was calculated and tested for statistical significance with a T-test. The results showed that the mean consistency score of new events with the original narrative was not significantly different from zero (T(14) = - 1.055, P = 0.309), which means the new events were perceived as neutral to the original narratives.

## RESULTS

Analyses of behavior revealed that the narratives were remembered well during the immediate free recall at the end of Day 1 (84 % accurate on average, Fig. 2A). On Day 2, in the MRI scanner, participants saw snapshots of a subset of previously experienced events (Old Narrative Events) and new events which could fit into those narratives (New Narrative Events). The presentation of Old and New Narrative Events was blocked, so that the participants first saw the 24 Old Narrative Events (six per narrative) several times, before they saw 6 New Narrative Events during the Updating Block that could relate to one of the families but had a neutral relationship with the two narratives about that family (which was defined based on control experiment 2, consistency task, details described in the methods section). The family which was updated with New Events was counterbalanced across participants. Thus, each New Narrative Event updated one of two competing narratives. During the Updating Block, participants also saw six new events which clearly did not fit with either family because they featured new characters (New Control Events). Subsequently, in a separate block, participants again saw the snapshots of the Old Narrative Events, which enabled us to measure the degree of reorganization of mnemonic networks as a consequence of memory updating.

**Fig. 2.**
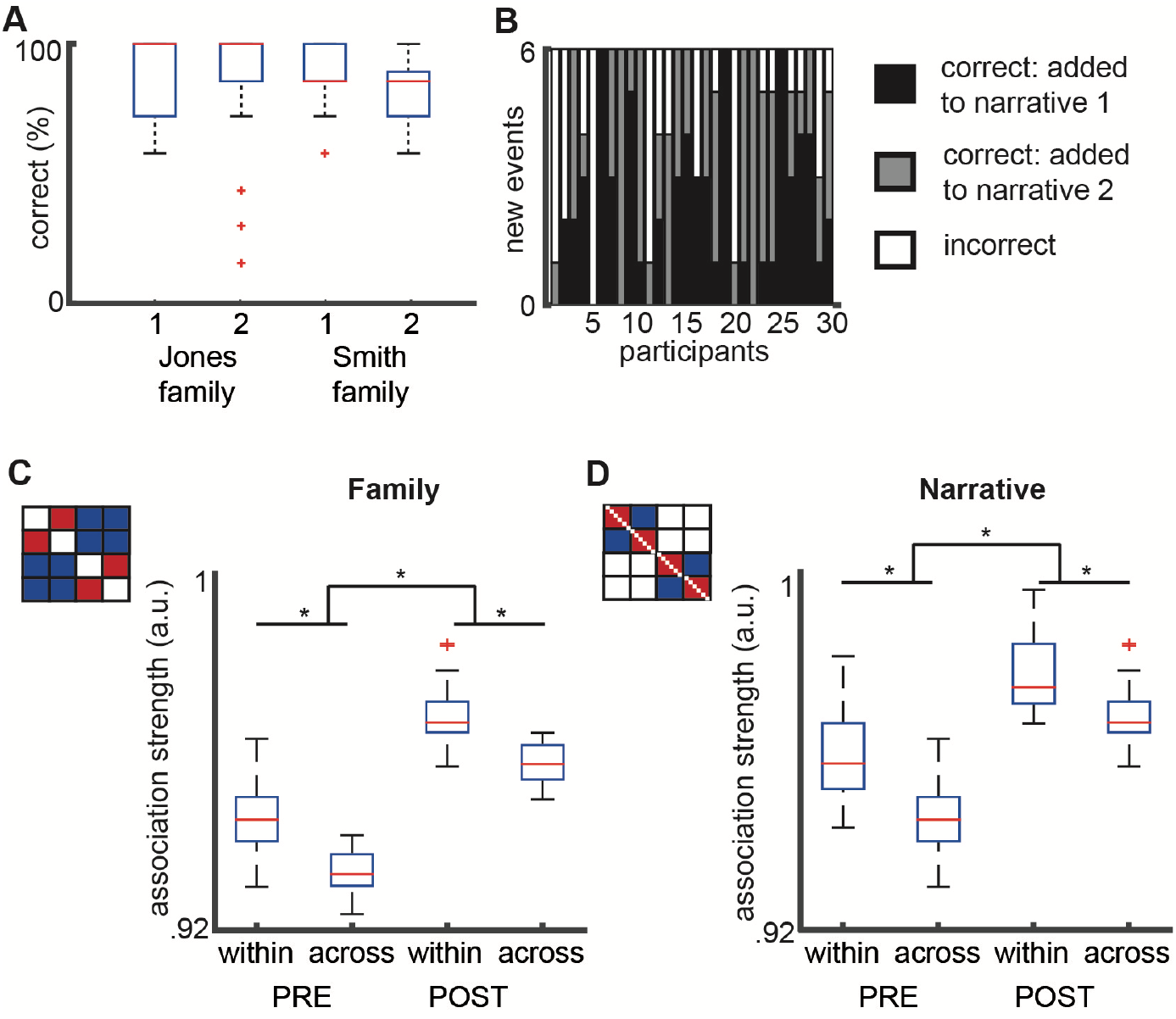
Behavioral results. **(A) Free recall test**. The participants (N=30) showed similar memory performance across all narratives (mean correct: 86%, 86%, 84%, and 80% for the four narratives separately; no significant difference in memory performance were observed between the narratives in a repeated-measures ANOVA: F(3,84) = 0.863, P = 0.435 Greenhouse-Geisser corrected). Results are visualized using boxplots with the median plus interquartile range, and the minimum and maximum values of the sample. The red crosses reflect potential outliers in that particular condition, but exclusion of those outliers does not influence the result. **(B) Updating performance**. For each participant separately, the number of new events that were positioned closest to narrative 1 of the updated family (black), narrative 2 of the updated family (gray), or to one of the narratives of the incorrect family (white). The new events were designed in such a way that they had a neutral relationship with the two narratives of the corresponding family – i.e. that they could be integrated into either narrative. On average, participants integrated 76.11% of New Narrative Events with the correct family, which was significantly above chance: T(29) = 4.59, P < 0.001. **(C) Integrating events from the same family**. Results from the pre and post scanning memory tests. The closer the participant has positioned the events of a certain category together, the higher the association strength (measured in artificial units, a.u.). Schematic shows the prediction matrix for the family-level-based grouping (i.e. Lower distance within-family, indicated in red, compared to across-family, indicated in blue, excluding within-narrative-pairs, indicated in white). Boxplots present association strengths averaged across updated and non-updated narratives since there was no interaction effect, i.e., no differential effect for updated versus non-updated narratives (boxplot visualizes the results with the median plus interquartile range, and min and max values of the sample). The red cross reflects a potential outlier in that particular condition, but exclusion of the outlier does not influence the result. **(D) Integrating events from the same narrative**. Results from the pre and post scanning memory tests. Boxplots present association strengths averaged across updated and non-updated narratives since there was no difference between results from updated and non-updated narratives (boxplot visualizes the results with the median plus interquartile range, and min and max values of the sample). The schematic shows the prediction for the narrative-level-based grouping (i.e. Lower distance within-narrative, indicated in red, compared to across-narrative, indicated in blue, excluding across-family-pairs, indicated in white). All experimental conditions in the analyses presented in this figure were measured in the same sample. The red cross reflects a potential outlier in that particular condition, but exclusion of the outlier does not influence the result.

### Behavioral results: Learning hierarchical event structures

Participants performed two event-configuration memory tests, one immediately before and one immediately after the MRI session. During these memory tests, participants arranged grayscale snapshots of the events (on a computer-screen) based on how these events “belong together”, which allowed us to quantify the structure of event networks of the narratives by calculating event-by-event distances between the events. Here, the participants were presented with snapshots of the events they had seen thus far (i.e. all Old Narrative Events were presented during the pre-MRI memory test, and all Old and New Narrative Events, as well as the Control Events, were presented in the post-MRI memory test), and had to arrange all these events at once on the computer-screen.

#### Integrating events from the same family

First, we tested whether participants grouped the events according to the two families. The results indeed revealed family-level-based groupings for both updated as well as non-updated in both pre and post memory tests (Fig. 2C and D; updated in pre memory test: one sample T-test, two-tailed, T(1,29) = 6.226, P = 0.000001; non-updated in pre memory test: one sample T-test, two-tailed, T(1,29) = 4.130, P = 0.000281; updated in post memory test: one sample T-test, two-tailed, T(1,29) = 4.368, P = 0.000146; non-updated in post memory test: one sample T-test, two-tailed, T(1,29) = 4.850, P = 0.00039).

#### Integrating events from the same narrative

Subsequently, we tested whether participants grouped the events according to the narratives. The results indeed revealed narrative-level-based grouping for updated as well as non-updated in both pre and post memory tests (Fig, 2C and D; updated narratives in pre memory test: one sample T-test, two-tailed, T(1,29) = 3.543, P = 0.00136; non-updated in pre memory test: one sample T-test, two-tailed, T(1,29) = 5.423, P = 0.000008; updated in post memory test: one sample T-test, two-tailed, T(1,29) = 3.887, P = 0.000543; non-updated in post memory test: one sample T-test, two-tailed, T(1,29) = 5.616, P = 0.000005).

#### Stronger integration after updating

A repeated-measures ANOVA with Time (pre memory test, post memory test), Integration-Level (Family, Narrative) and Updated-Family (Yes, No) as within-subject factors showed a significant main effect of time, with stronger family and narrative integration of events in the post memory test compared to pre memory test (F(1,29) = 7.102, P = 0.012). This suggests that the MRI session during which Old as well as New Narrative Events were shown further integrated the events of the narratives.

### The Hippocampus integrates New Narrative Events

We next examined the neuroimaging data. First, we investigated possible integration of the New Narrative Events by performing a contrast which predicted high similarity between the six New Narrative Events and low similarity between the six New Control Events. As expected, the results indeed showed higher neural similarity in the hippocampus between the New Narrative Events compared to the neural similarity between the New Control Events (small volume corrected for the entire left and right hippocampus; Fig. 3A and Table 1; peak searchlight in hippocampus, MNI coordinates X= −30, Y= −14, Z= −20; T = 4.17). Control analyses showed that the effect was not caused by difference in amplitude of the BOLD signal or by differences in low-level visual features (see Methods for more details). To examine whether this increase in neural similarity could be explained in terms of overlapping features, in this case characters, we performed a control analysis that predicted high similarity between New Narrative Events which featured the same characters and low similarity for New Narrative Events with different characters (New Control Events were not used in this analysis as each event contained different characters). Increased similarity for events with overlapping features was observed in the primary visual cortex (peak searchlight, MNI X = 12, Y = −86 and Z = 0, T = 5.4), but not in the hippocampus, suggesting that the differentiation between New Narrative and Control Events was not driven by common characters across Narrative events.

**Fig. 3.**
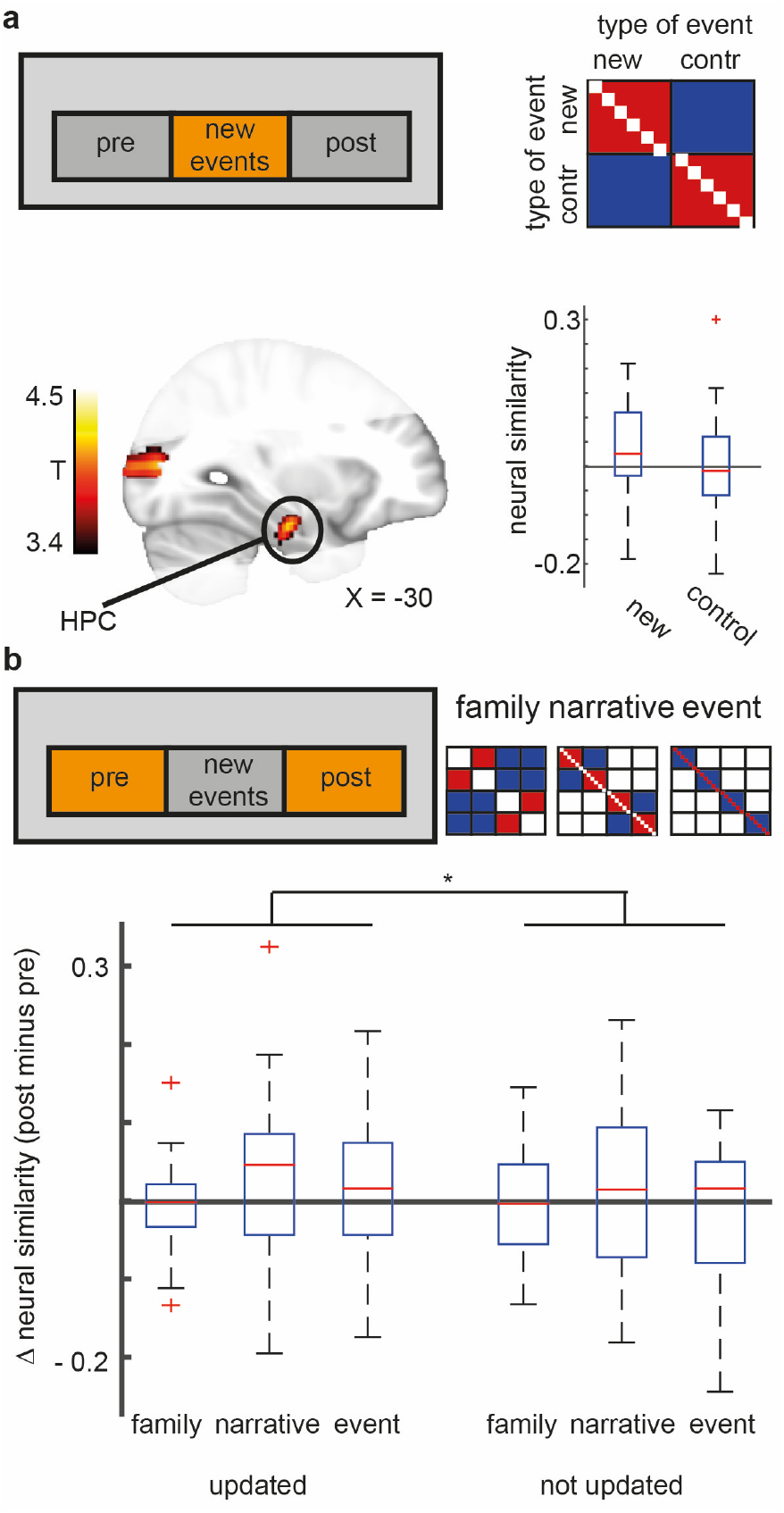
Representation of new narrative events and strengthening associations between old narrative events after updating. **(A) Representation of new events**. Participants (N=30) showed a higher neural similarity between the six New Narrative Events than between the six Control Events in the hippocampus (one sample T-test, small volume corrected for the entire left and right hippocampus, peak voxel MNI coordinates X = −30, Y = −14, Z = −20; T = 4.17). Top right shows the prediction matrix we used (red means high predicted neural similarity, blue means low predicted neural similarity, white means that this cell was excluded from this analysis). Brain figures indicate the field-of-view of the MR sequence we used for this experiment. Statistical significance of the hippocampus is based on a small volume correction, because this region was our a priori region of interest. Brain figures displayed in the figure are P < .001 uncorrected for display purposes. Whole brain corrected results are reported in table 1 (P < 0.05 FDR corrected). Additionally, a boxplot visualization (median plus interquartile range, and min and max values of the sample) of this effect for the peak searchlight in the hippocampus (center of peak searchlight MNI coordinates X = −30, Y = −14, Z = −20) is shown at the bottom right of panel A. The red cross reflects a potential outlier in that particular condition, but exclusion of the outlier does not influence the result. **(B) Strengthening associations between old narrative events after memory updating**. We ran three post hoc analyses on the pre and post blocks to test the influence of updating on the original narrative representations in the anterior hippocampus (corresponding to location of effect in panel A). We tested the three levels of the narrative hierarchy: [1] higher similarity between events of the same family compared to across family, [2] higher similarity between events of the same narrative compared to across narratives, and [3] higher similarity within compared to across events. We used prediction matrices as shown in the figure, with red means high predicted neural similarity, blue means low predicted neural similarity, white means that this cell was excluded from this analysis. To examine what information about the narrative hierarchy is present in the voxels that represented the New Narrative Events, we extracted the peak searchlight of the hippocampus effect from panel A. The results tested with a repeated-measures ANOVA with Session (Pre, Post), Updated (Yes, No) and Hierarchical-level groupings (Family, Narrative, Event) as within-subject factors revealed a significant Updated x Session interaction effect (F(1,29) = 4.548, P = 0.042). All experimental conditions in the analyses presented in this figure were measured in the same sample. Boxplot visualizes the results with the median plus interquartile range, and the minimum and maximum values of the sample. The red crosses reflect potential outliers in that particular condition, but exclusion of those outliers does not influence the result.

**Table 1.** Summary table of representational similarity results: Representation of new narrative events. Statistics are reported for peak voxels surviving a whole field-of-view FDR-correction (P < .05).

|  | MNI coordinates |  |  |  |
| --- | --- | --- | --- | --- |
| Region | X | Y | Z | T-value |
| Hippocampus<sup>1,2,3</sup> (extending into parahippocampal gyrus<sup>1</sup>) | -30 | -14 | -20 | 4.17 |
| Frontal inf<sup>3</sup> | -40 | 16 | 20 | 4.50 |
| Occipital mid<sup>1,2,3</sup> | -36 | -80 | 10 | 4.40 |
| Frontal med orb<sup>1</sup> | 4 | 68 | -6 | 4.30 |
| Cerebellum<sup>3</sup> | -2 | -72 | -36 | 4.29 |
| Rectus | 0 | 46 | -20 | 3.88 |
| Temporal mid<sup>1,3</sup> | 40 | -60 | 10 | 3.76 |
| Fusiform gyrus<sup>2</sup> | -26 | -46 | -16 | 3.66 |
| Temporal pole<sup>1</sup> | -20 | 14 | -32 | 3.46 |
| Frontal sup<sup>1,3</sup> | -14 | 72 | 8 | 3.44 |
| <sup>1</sup> Part of the autobiographical memory network (see Spreng et al., 2008).<br><sup>2</sup> Showed narrative representation in Milivojevic et al., 2016.<br><sup>3</sup> Showed narrative representation in Milivojevic et al., 2015. |  |  |  |  |

### Better integration of Old Narratives Events due to updating in the hippocampus

Next, we tested whether the configuration of pre-existing hippocampal event networks changed as a consequence of introducing these New Narrative Events. We predicted that any representational change would be restricted to those Old Narrative Events which feature the same family as the New Narrative Events. Do the same voxels that carry information about the New Narrative Events (see Fig 3A) also carry information about the modification of Old Narrative Event representations? We tested whether the hippocampus (search sphere defined around the peak searchlight that showed a significant effect in the RSA searchlight shown in Fig. 3A) further integrated the Old Narrative Events according to the three levels (i.e., event, narrative, and family). Thus, we tested 3 predictions for which we used: [1] a contrast that predicted higher neural similarity within, compared to across, events (event representation), [2] a contrast that predicted higher neural similarity for within-narrative event pairs compared to across-narrative event pairs (narrative representation), and [3] a contrast that predicted higher neural similarity for within-family event pairs compared to across-family event pairs (family representation). A repeated-measures ANOVA with Session (Pre, Post), Updating (Yes, No) and Level (Family, Narrative, Event) as within-subject factors revealed a significant Updating x Session interaction (F(1,29) = 4.548, P = 0.042), with selective increases in similarity for the updated family after updating (relative to the non-updated narratives and the pre block, see Figure 3B). These results suggest that updating with New Narrative Events evoked sharpening of narrative representations in the hippocampus for specifically those narratives that were updated with new events.

### Hippocampus integrates New Narrative Events with Old Narrative Events

Since the task required participants to actively place the New Narrative Events at a specific moment within an Old Narrative, we expected high neural similarity of New with Old Narrative Events, selectively for that family which was featured in the new events (Fig. 4A). Thus, we asked whether the hippocampus showed high neural similarity of New with Old Narrative Events, by performing a contrast that predicted high similarity between New Narrative Events and Old Narrative Events from the updated family, and low similarity between New Narrative Events and the Control Events (i.e. Old Narrative Events from the non-updated family and New Control Events). We compared neural similarity between New Narrative Events presented in the updating block with Old Narrative Events presented in the post block. The results indeed revealed higher neural similarity in the hippocampus between New and Old Narrative Events of the updated family, compared to Control Events and non-updated family (small volume corrected for the entire left and right hippocampus; Fig. 4B and C and Table 2; center of peak searchlights in hippocampus, MNI coordinates X= −26, Y= −36, Z= −6; T=4.47 and X= −16, Y= −40, Z= 6; T=4.01).

**Fig. 4.**
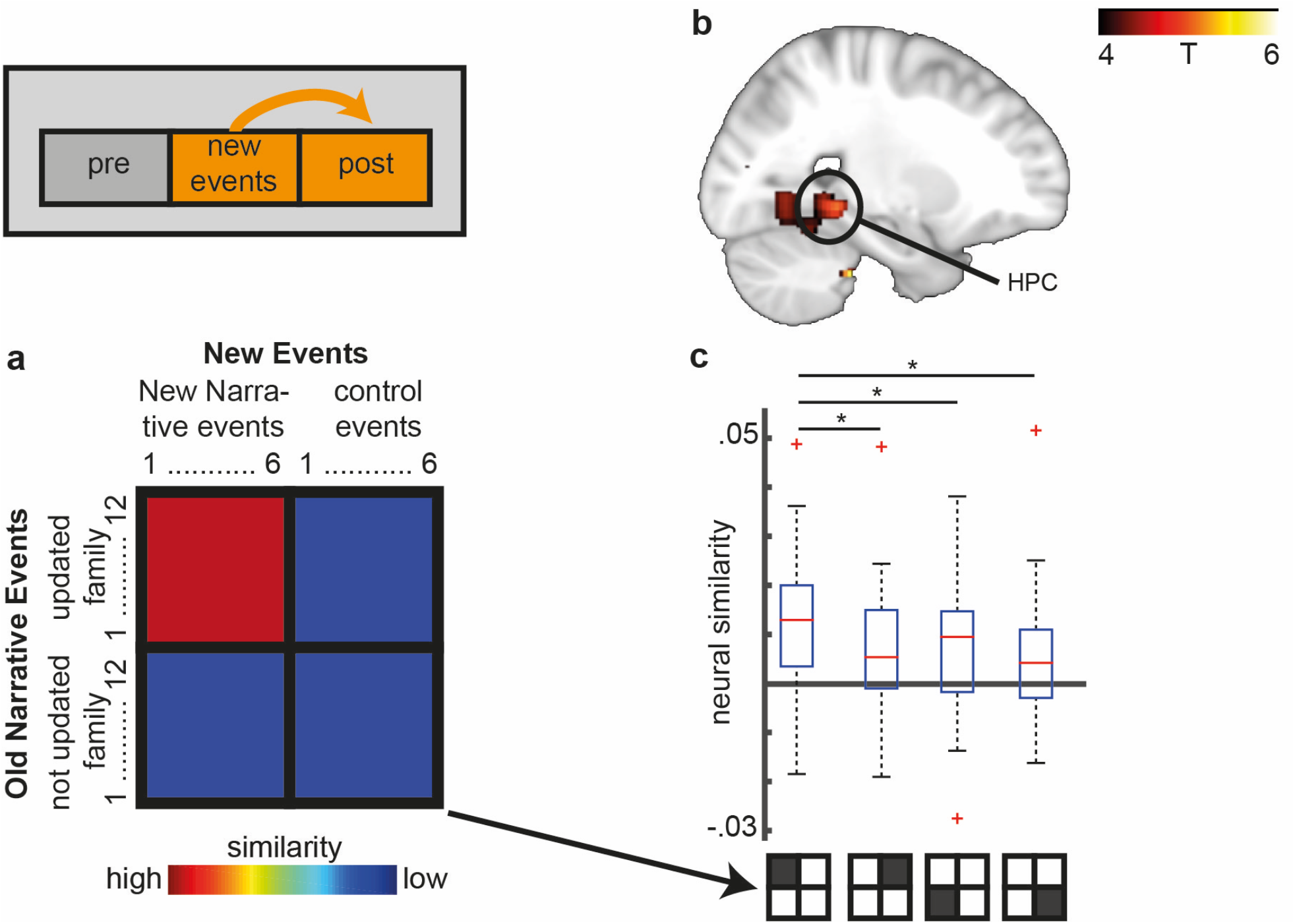
Integration of Old and New Narrative Events. **(A) Prediction matrix**. We ran an analysis (N=30) which predicted higher neural similarity between the six new events (in the updating block) with their twelve corresponding original Narrative Events (in the post block), relative to the controls family and Control Events (see panel C for more details). Red means high predicted neural similarity, blue means low predicted neural similarity. **(B) Integration of new events with their corresponding narratives**. This analysis revealed higher neural similarity between New Narrative Events (presented in the updating block) and their corresponding Old Narrative Events (presented in the post block), relative to controls, in the hippocampus (small-volume corrected for the hippocampus; one sample T-test, peak searchlight MNI coordinates X = - 26, Y = - 36, Z = - 6; T = 4.47). Table 2 lists other regions that showed a significant effect. For the main result presented here, new narrative events from the updating block were compared with corresponding old Narrative Events from the post block. The results for increased similarity of new narrative events with the pre block is presented in Table 2. **(C) Visualisation of effect shown in panel B**. In order to gain a better understanding of the integration effect presented in panel B, we here present the neural similarity of each of the quadrants of the prediction matrix separately (new events to updated family, Control Events from updated family, new events to not updated family, Control Events to not updated family). Post-hoc two-tailed paired-samples T-tests showed that the higher neural similarity is indeed driven by the high similarity between new events and their corresponding narratives, relative to the other three quadrants of the prediction matrix of panel A (from left to right: T(1,29) = 2.397, P = 0.023, T(1,29) = 2.246, P = 0.033, T(1,29) = 2.413, P = 0.022). Boxplot visualizes the results with the median plus interquartile range, and min and max values of the sample. All experimental conditions in the analyses presented in this figure were measured in the same sample. The red crosses reflect potential outliers in that particular condition, but exclusion of those outliers does not influence the result.

**Table 2.** Summary table of representational similarity results: integration of New with Old Narrative Events. Statistics are reported for peak voxels surviving a whole field-of-view FDR-correction (P < .05).

|  | MNI coordinates |  |  |  |
| --- | --- | --- | --- | --- |
| Region | X | Y | Z | T-value |
| **Similarity of new narrative events with old narrative events of pre and post blocks together** |  |  |  |  |
| Occipital mid<sup>1,2,3</sup> | 42 | -86 | 22 | 4.82 |
| Temporal mid<sup>1,3</sup> | 54 | -64 | 24 | 4.28 |
| Hippocampus<sup>1,2,3</sup> (extending into parahippocampal gyrus<sup>1</sup> and fusiform gyrus<sup>2</sup>) | -24 | -40 | -4 | 5.81 |
| Temporal inf<sup>1</sup> | 58 | -60 | -22 | 4.82 |
| Cuneus<sup>1</sup> | 20 | -70 | 24 | 4.21 |
| Frontal inf<sup>3</sup> | 54 | 10 | 30 | 3.76 |
| Frontal sup<sup>1,3</sup> | -2 | 54 | 30 | 3.16 |
| Cerebellum<sup>3</sup> (extending into lingual gyrus) | 4 | -62 | -4 | 6.36 |
| **Similarity of new narrative events with old narrative events of pre block only** |  |  |  |  |
| Occipital mid<sup>1,2,3</sup> | -48 | -70 | 0 | 4.76 |
| Temporal mid<sup>1,3</sup> | -66 | -42 | 2 | 4.73 |
| Fusiform gyrus<sup>2</sup> | 44 | -34 | -24 | 4.38 |
| Cerebellum<sup>3</sup> | 4 | -66 | -4 | 5.32 |
| Frontal sup<sup>1,3</sup> | -4 | 54 | 32 | 3.85 |
| Thalamus<sup>1</sup> | -12 | -8 | 14 | 3.30 |
| Frontal inf<sup>3</sup> | 56 | 12 | 30 | 3.16 |
| **Similarity of new narrative events with old narrative events of post block only** |  |  |  |  |
| Occipital mid<sup>1,2,3</sup> | -38 | -74 | 2 | 5.50 |
| Temporal mid<sup>1,3</sup> | 50 | -66 | 8 | 4.56 |
| Fusiform gyrus<sup>2</sup> | 34 | -46 | -6 | 4.00 |
| Hippocampus<sup>1,2,3</sup> | -26 | -36 | -6 | 4.47 |
| Hippocampus<sup>1,2,3</sup> | -16 | -40 | 6 | 4.01 |
| Temporal inf<sup>1</sup> | 60 | -60 | -20 | 4.07 |
| Cuneus<sup>1</sup> | 20 | -68 | 24 | 4.15 |
| Cerebellum<sup>3</sup> | -26 | -34 | -38 | 5.59 |
| Insula | -34 | 18 | 14 | 3.75 |
| Frontal sup<sup>1,3</sup> | -18 | 64 | 8 | 3.61 |
| <sup>1</sup> Part of the autobiographical memory network (see Spreng et al., 2008). |  |  |  |  |
| <sup>2</sup> Showed narrative representation in Milivojevic et al., 2016. |  |  |  |  |
| <sup>3</sup> Showed narrative representation in Milivojevic et al., 2015. |  |  |  |  |

Control analyses showed that the effect was not caused by difference in amplitude of the BOLD signal, nor by differences in low-level visual features (see Methods).

### No evidence for family level integration of events

Next, we investigated integration of events at the family level. We predicted that any representational change would be restricted to those Old Narrative Events which feature the same family as the New Narrative Events. Thus, we tested an interaction contrast that predicted higher neural similarity for within-family events compared to across-family events in the post compared with the pre-updating block. A searchlight analysis revealed no significant effects anywhere, including the mPFC and hippocampus, even at a liberal P < 0.001 threshold uncorrected for multiple comparisons. A possible explanation for the absence of this predicted effect could be that the task specifically required participants to place each new events at a specific timepoint within the timeline of a single narrative which might have caused the two narratives from that family to be competing, and therefore not integrated. Additionally, it suggests that narratives are preferably used as a context for organizing episodic memories rather than a more overarching aspect (like Family in this design).

## DISCUSSION

We examined the reconfiguration of event networks in the hippocampus by presenting new events that could update either one of two competing event networks. We identified three hippocampal mechanisms for updating event networks. The hippocampus was sensitive for a network representation of new experiences during updating. Updating caused further integration of related experiences from the past in behavior and in the pattern of hippocampal activity. The hippocampal representations of these new experiences were integrated with the hippocampal representations of related experiences from the past.

The results revealed increased neural similarity between all new events in the anterior hippocampus. This finding suggests a key role for the anterior hippocampus in the creation of event networks, which is in line with earlier work suggesting that memory hierarchies are represented along the hippocampal long axis with networks at a large scale represented anteriorly (Komorowski et al., 2013; McKenzie et al., 2014; Collin et al., 2015; Brunec et al., 2018). These findings are in line with evidence that showed that the anterior hippocampus is particularly involved in memory integration and inference (Collin et al., 2015; Schlichting et al., 2015; Deuker et al., 2016). These results also dovetail with our previous studies where we discovered increased hippocampal pattern similarity for related events (Collin et al., 2015; Milivojevic et al., 2015) and for events from the same narrative (Milivojevic et al., 2016), which we interpreted as indicative of the formation of an event network in the hippocampus. This is consistent with prominent theories suggesting that the hippocampus creates a memory space (Eichenbaum et al., 1999) or cognitive map (O’Keefe and Nadel, 1978) of related events.

Besides a network of new events, the anterior hippocampus also strengthened the event networks which consisted of all events of earlier experiences related through individual narratives. Importantly, this was only evident after narratives had been updated and was specific to the updated narratives, which suggests it was triggered by the updating manipulation. This is in accordance with animal reconsolidation studies suggesting that additional learning strengthens established memories (Dudai and Eisenberg, 2004; Lee, 2008). Strengthening of old experiences in anterior hippocampus after updating might suggest that updating was accompanied by reactivation of relevant context or experiences in the anterior hippocampus, which then remained online during the post-updating block. Largely in line with these imaging findings, the updating manipulation also triggered strengthening of earlier experiences in the pattern of behavior. In contrast to the imaging findings, however, the behavioral results suggest strengthening of earlier experiences for both narratives of the updated and also the non-updated family. It is difficult to reconcile these differences in neural similarity and behavioral results, as they appear conflicting. It is possible that during the behavioral task, which requires simultaneous arrangement of multiple events along the screen, strengthening of relations between events of the updated family also led the participants to place the remaining events (i.e. of non-updated family) closer together.

Previous findings revealed posterior hippocampus involvement in representing distinct events (Schlichting et al., 2015), and in changing the interrelationships between certain events, i.e. flexibly ‘re-linking’ events after new information was introduced about how these events belonged together (Collin et al., 2015; Milivojevic et al., 2015). The increased neural similarity of new with old events in the posterior hippocampus found in this study is indicative of the engagement of the posterior hippocampus in the flexible integration of new information with previous experiences. This result indicates that posterior hippocampus might not represent the networks of new or earlier experiences themselves, but rather associations across separate networks, and thus representing information about specific, detailed, relationships between old and new experiences. This posterior hippocampal involvement in remembering the precise relationship between new and old experiences might relate to a phenomenon referred to as remapping, which is the formation of distinct representations of hippocampal place cells after changes in input to the hippocampus (Colgin et al., 2008; also Steemers et al., 2016 and Julian and Doeller, 2021, for fMRI evidence for hippocampal remapping). The posterior hippocampus might aid in creating another representation after a change in hippocampal input that includes new information, which is in line with the view that pre-existing structures can be modified to accommodate new information (Piaget, 1929). Together, the data suggests that three critical components of updating episodic memories in the hippocampus, i.e. 1) memory representation of new events, 2) integration of new with related old events, and 3) sharpening of old events, underlie updating of memory networks when new information comes to light.

The results could suggest that the anterior hippocampus maintains the information about the source of the memories: it maintains the information about which events belong to new experiences and which belong to old experiences by representing these events in two separate networks in parallel. In line with this, the anterior hippocampus is indeed known to be important for representing context (Collin et al., 2015; Brunec et al., 2018; Guo and Yang, 2020). The fact that anterior hippocampus represents networks of old and new experiences separately could be interpreted as a way to keep track of the temporal context of experiences. This notion that the anterior hippocampus also keeps track of the temporal relationship between related events is in line with multiple studies showing evidence that anterior temporal lobe represents the temporal context of events (Ezzyat and Davachi, 2014; Deuker et al., 2016; Lositsky et al., 2016). In our study, the anterior hippocampus might simultaneously keep multiple networks of multi-event experiences that are encoded at different moments in time as separate networks, because it might be tracking relations between events from the same temporal context (Howard and Kahana, 2002; Staresina and Davachi, 2009; Hsieh et al., 2014; Deuker et al., 2016; Ranganath and Hsieh, 2016; Cohn-sheehy and Ranganath, 2017; Lee et al., 2020; Bellmund et al., 2021). The role of the posterior hippocampus might then be to provide additional detailed information about how these experiences fit together. Integrating events from a single experience and simultaneously pattern-separating multiple related experiences (Yassa and Stark, 2011) is critical for representing hierarchical structures (Kumaran, 2013; McKenzie et al., 2014; Brod et al., 2015). The differential involvement of anterior and posterior hippocampus might enable the hippocampus to later on retrieve a coherent memory of all events of a single experience without catastrophic interference (Hasselmo, 2017), which is in line with the known functions of pattern separation and pattern completion in the hippocampus (Leutgeb and Leutgeb, 2007; Yassa and Stark, 2011; Rolls, 2013; Richards and Frankland, 2017). Additional work is necessary to draw conclusions about the nature of putative parallel mnemonic codes in the hippocampus. It should also be noted that in some cases it is advantageous for memory traces to be forgotten for the purpose of generalization or formation of unified narrative representations. It is likely that the separate representations may merge once the updated memories undergo a longer period of consolidation (Richards and Frankland, 2017).

We found no evidence of updating in the mPFC. It is possible that the lack of the mPFC involvement reflects the complex task demands for this experiment (i.e., the participants were instructed to insert each new event at a specific position), whereby the events need to be flexibly recombined into pre-existing narratives – a process which critically depends on the hippocampus (Wikenheiser and Schoenbaum, 2016; Gilboa and Marlatte, 2017). Alternatively, it is conceivable that these complex structures required more time before it is consolidated into schematic representations in the mPFC, and future studies could examine whether allowing additional time for consolidation would result in strengthening of mPFC representations after updating.

The increases in neural similarity as outlined here can be interpreted as the creation of integrated representations. Alternatively, the increases in neural similarity could reflect directly associating representations of different events or indirectly integrating the representations of different events by binding them all to one common representation, however, as discussed previously (Milivojevic et al., 2015), these alternative models are less likely. Additionally, the neural similarity increases could reflect common processes engaged for narrative events or the activation of shared features while watching narrative events. However, also control events shared features with other control events, while at the same time critically lacking a shared prior experience, thereby ensuring that similar associative processes were engaged for both control events and narrative events, but only narrative events required updating of pre-existing neural representations. We conclude that the increases in neural similarity as outlined here are most likely caused by the creation of shared representations after linking events together.

In conclusion, we examined whether updating of narrative-based memory networks relies on hippocampo-cortical mechanisms. Here we present three hippocampal mechanisms for updating event networks in memory: [1] integration of new events, [2] integrating these new events with earlier events, and [3] better integration of earlier events after updating. These results suggest that memories are organized into dynamic networks based on narrative contexts. With these networks the hippocampus can keep track of how all our experiences over time relate to each other without losing the ability to remember the contextual specifics of when those events were experienced.

## ACKNOWLEDGMENT

This work was supported by the Netherlands Organisation for Scientific Research (NWO-Vidi 452-12-009). CFD’s research is further supported by the Max Planck Society; the European Research Council (ERC-CoG GEOCOG 724836); the Kavli Foundation, the Jebsen Foundation, the Centre of Excellence scheme of the Research Council of Norway – Centre for Neural Computation, The Egil and Pauline Braathen and Fred Kavli Centre for Cortical Microcircuits, the National Infrastructure scheme of the Research Council of Norway – NORBRAIN. SHPC was further supported by the Netherlands Organisation for Scientific Research (NWO-Rubicon grant 446-17-009). Correspondence should be addressed to SHPC or CFD. The authors would like to thank JLS Bellmund and KA Norman for helpful feedback on an earlier version of this manuscript.

## CONFLICT OF INTEREST

The authors declare no competing financial interests.

## Notes

### Competing Interest Statement

The authors have declared no competing interest.

